# When the inner clock fades: Interoceptive decline and consolidation of phase resetting in cortical rhythms by cardiac events underlie healthy lifespan ageing

**DOI:** 10.1101/2024.05.31.596844

**Authors:** Kirti Saluja, Dipanjan Roy, Arpan Banerjee

## Abstract

The brain continuously tracks signals from the body, including the heartbeat, which influence internal states such as the perception of time, well-known to accelerate with age and disrupted in disorders such as Parkinson’s disease and dementia. The present study examined whether the representation of cardiac events onto spontaneous neural oscillations maps to regions responsible for processing time-perception, show age-related differences, and uncovered the mechanisms that may underlie any such reorganization. From a large cohort (N = 620), cortical heartbeat-evoked responses (HERs) were characterized and their sources were pinned to frontotemporal regions. Phase-based analyses demonstrated that cardiac signals affect phase of ongoing neural oscillations in the theta frequency band, rather than altering overall power, with this effect consolidating in older adults. Advanced statistical analysis further indicated that these changes are primarily driven by enhanced bottom-up heart-to-brain influences, revealing that altered interoceptive signaling can shape the age-related changes in time perception.

## Introduction

Time perception is a core aspect of human cognition, yet it undergoes systematic alterations across the lifespan. A well-established phenomenon is the subjective acceleration of time with advancing age^1^, a pattern also observed in neurodegenerative conditions such as Parkinson’s disease ^2^ and dementia^3^. Emerging evidence suggests that temporal perception is closely intertwined with internal bodily rhythms, particularly cardiac activity, indicating that heart may serve as a dynamic internal clock that helps structure our experience of time. Cardiac signals, specifically the systolic and diastolic phases of the heartbeat, are increasingly recognized as contributors to temporal processing in the brain^4^. However, the mechanisms through which these heart-brain interactions shape our perception of time, and how these mechanisms evolve with age – remain poorly understood. One promising avenue for investigating this interaction involves heartbeat-evoked responses (HERs), which reflects the brain’s response to afferent cardiac signals^5–9^. These signals are transmitted via the vagus nerve and dorsal root ganglia to brainstorm and thalamus, and then to the higher-order cortical areas including the hypothalamus, amygdala, anterior cingulate cortex, insula, and orbitofrontal cortex^10^. HERs have been implicated in a wide range of cognitive and affective processes, including arousal^11^, predictive coding^12,13^, interoceptive accuracy in emotional tasks^14,15^, heart disease^16^, somatosensory perception^17^, attention^18^, consciousness^19^, mental disorders^20–22^ and importantly, aging^23^ and time perception^4^.

Despite this progress, except this study^8^, all majority of HER studies have focused linking task performance with HERs – for instance, by examining whether specific phases of cardiac cycle enhance perceptual or cognitive outcomes. This task-based framing, however, limits our understanding of truly endogenous, spontaneous neural responses to the heartbeat, which can predict task performance even during pre-stimulus intervals. Furthermore, it proposed that heartbeats can induce phase synchronization within the theta frequency band across cortical regions during rest, forming what they termed the heart-induced network (HIN) further connected to emotion regulation and interoception. To uncover the neural foundations of time perception as it naturally evolves across the lifespan, a focus on resting-state HERs becomes essential.

Additionally, another study ^23^ reported age-associated alterations in the HER amplitude is correlated with the performance in the metacognitive judgement task. Even though this study is pertinent given ageing affects both structural and functional aspects of the brain^24^, the study is constricted with datasets focused on extreme age groups, and few examine how HER-related networks are reorganized across the adult lifespan. Furthermore, the network-level mechanisms by which cortical representations of cardiac signals consolidate or decline with age remain largely unexplored. A more granular, lifespan-continuous approach, examining how HERs manifest across age in resting-state brain activity, could help clarify the underlying mechanisms that contribute to the subjective speeding of time in aging. Such findings may also offer biomarkers for pathological aging in disorders like Alzheimer’s and Parkinson’s disease.

Based on this foundation, the present study asks: Does the resting brain carry age-dependent signatures of time perception, as reflected in its representation of cardiac signals? Subsequently, a hypothesis whether brain regions known to support time perception in psychophysical tasks also represent cardiac events during rest is tested, following which it revealed that this representation becomes more consolidated with age - a possible substrate for the age-related acceleration in subjective time perception. Resting state magnetoencephalography (MEG) data from the Cambridge Centre for Ageing and Neuroscience (Cam-CAN) cohort (N = 620, aged 18–88 years) were utilized from which HERs are extracted, time-locked to the R-peaks of the electrocardiogram (ECG). Inter-trial phase coherence (ITPC) was computed to assess whether cardiac events induce phase-resetting in cortical oscillations—a mechanism proposed to encode temporal events. Further, to explore the causal nature of heart-brain interactions, particularly within high-frequency neural oscillations and heart rate variability, Granger causality^25–27^ was used to uncover directional influences in heart-brain communication. Together, these analyses aim to provide a deeper understanding of the circuit-level principles underlying time perception across the human lifespan.

## Results

### Physiological metrics of cardiovascular health across lifespan ageing

The pipeline presented in Figure 1 was used to analyse the electrocardiographic (ECG) and magnetoencephalographic (MEG) data from the Cam-CAN cohort ^28^, comprising 620 participants aged 18 to 88 years.

**Figure 1:**
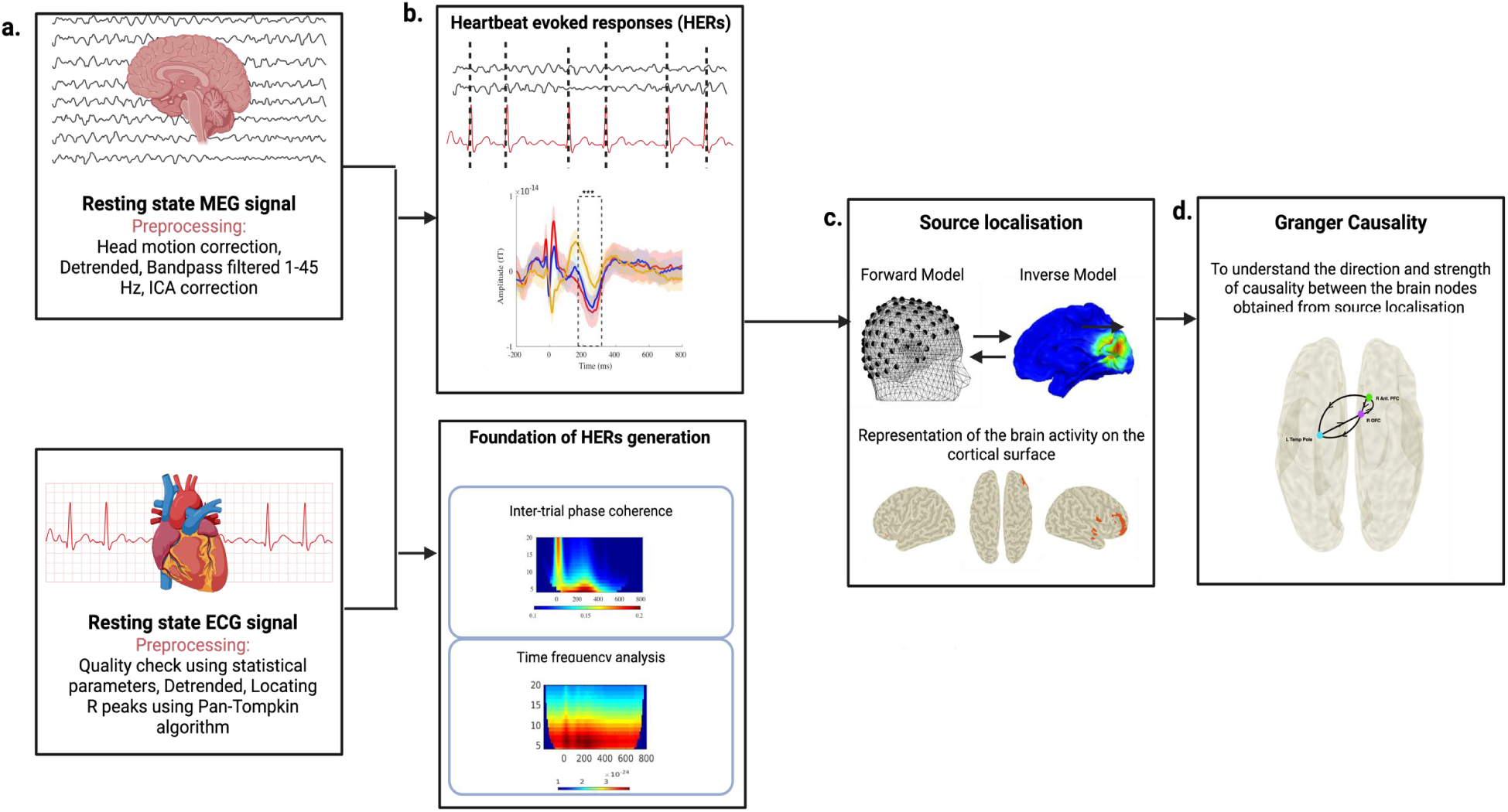
Overview of the Data Pre-processing and Analysis Pipeline. (a) Simultaneous MEG-ECG recordings undergo a preprocessing pipeline including detrending, bandpass filter and artifact removal using Independent Component Analysis (ICA). R peaks are detected from the electrocardiogram (ECG) waveform using Pan-Tompkin algorithm, serving as a trigger point to compute heartbeat evoked responses (HERs) (b) HERs are computed across all the age-groups (young, middle and old). Additionally, wavelet analysis is employed to investigate the mechanisms underlying HERs generation (c) Source localisation using eLORETA identifies cortical sources underlying HERs (d) Finally, Granger causality maps casual functional relationships among cortical sources of HERs.

To assess the cardiovascular health, electrocardiographic parameters were utilized including inter-beat interval (IBI), heart rate variability (HRV), heart rate (HR) and blood pressure (BP) indicators such as systolic and diastolic pressure (Figure 2.a).

**Figure 2:**
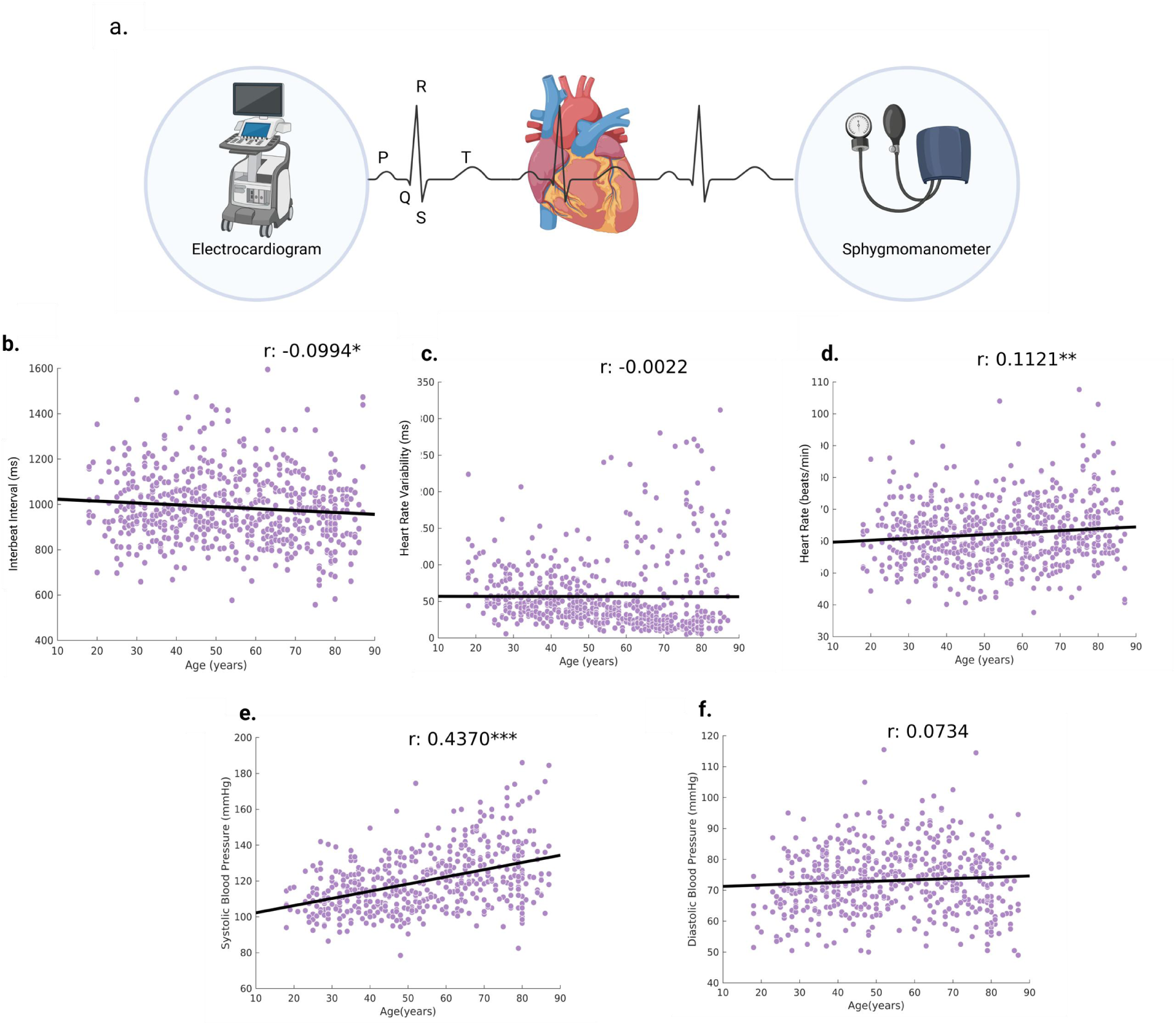
Cardiovascular measures across age. a. Methods used to capture cardiac activity includes electrocardiogram (ECG) and sphygmomanometer which measures cardiovascular activities such as electric activity of the cardiac muscles and blood pressure respectively. Cardiovascular parameters include – Inter beat interval (IBI), Heart rate variability (HRV), Heart rate (HR), Systolic and Diastolic blood pressure b. IBI significantly decreases (r = -0.0994; p = 0.0133) c. HRV doesn’t change (r = -0.0022; p = 0.9573) d.HR significantly increases (r = 0.1121; p=0.0052) e. Systolic blood pressure significantly increases (r = 0.4370; p<0.001) and f. Diastolic blood pressure remains unchanged (r=0.0734; p= 0.0910) across age.

These parameters were derived from ECG time series and blood pressure scores provided by Cam-CAN. IBI significantly decreases (r = -0.0994, p = 0.0133; Figure 2.b) and HR significantly increases across age (r = 0.1121, p = 0.0052; Figure 2.d) (p < 0.05). However, HRV does not decrease significantly across age (r = -0.0022, p = 0.9573; Figure 2.c). In case of the trend of blood pressure scores across age, systolic pressure was significantly increasing (r = 0.4370, p<0.001; Figure 2.e) whereas diastolic pressure was significantly increasing (r = 0.0734, p = 0.0910; Figure 2.f). Refer to the supplementary table S1 for additional details regarding the model.

### Heartbeat evoked responses (HERs) across lifespan

Simultaneous recordings of ECG and MEG were considered to create epochs of cardiac cycle comprising a QRS waveforms of 1 s duration, with first 200 ms representing pre-R peak period followed by 800ms post R peak (Figure 3.a). The epochs were averaged to compute the HERs channel-by-channel. The global HERs calculated for three age groups i.e., young (18-40 years), middle (41-60 years), and old (61-87 years) (Figure 3.a), as well as for genders groups i.e., male and female (Figure 3.e). The main effect of age on HERs (F (2,619) = 9.47, p = 0.0001) was significant while the effect of gender became non-significant (F(1,619) = 0, p =0.9631) in the early phase (180-320ms post R peak) window. Furthermore, an interaction of both factors were found to be non-significant (F (2,619) = 0.33, p = 0.7213). Refer to supplementary table S2 for further details. To validate this result, the HERs were calculated with respect to the shuffled/ surrogate ECG time series (see Methods for detail) and, no time-locked activity was observed relative to the non-R peak points (Figure 3.c).

**Figure 3:**
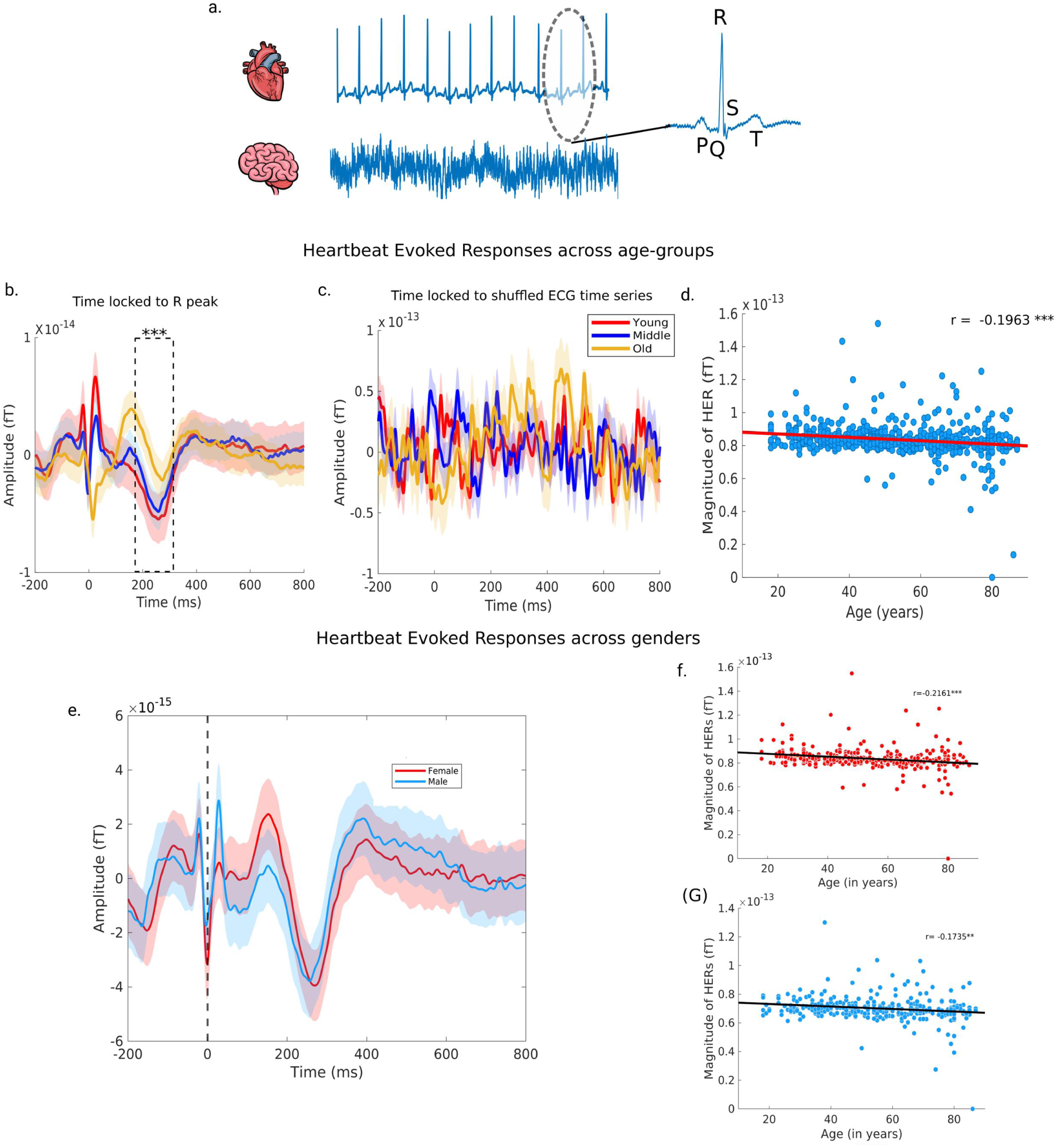
Heartbeat evoked responses (HERs) across age and gender. a. Method: Simultaneous MEG-ECG recording: using R peak of the ECG waveform as the time points to epoch neural time series to obtain the HERs (b) Group wise analysis of HERs obtained from the time locked MEG activity to R peak of the ECG waveform. Statistical test conducted indicates that the HERs differences across the age groups are significant shown by the asterisks (F(2,619) = 9.48; p<0.001) c. Group wise analysis of HERs obtained from time locked activity to the shuffle ECG time series (sanity check) d. Linear regression conducted between magnitude (obtained from peak to peak subtraction) of HERs and age (r= -0.1963; p<0.001) e. shows the group-wise analysis of HERs i.e., time locked MEG activity to the R peak (represented by the dotted line) of the ECG waveform for two gender (male and female). Statistical test conducted indicated that there is no significant difference between the two groups (F (1,619) = 0.02; p = 0.8801). The shaded area shows the standard deviation. Linear regression conducted magnitude (obtained from peak to peak subtraction) of HERs across age in f. Females (r= -0.2161; p<0.001) and g. Males (r = - 0.1735; p<0.01).

Further, linear regression analysis shows that the HERs amplitude extracted from the window of 180-320ms post R peak of the ECG waveform significantly decreases across age (r=-0.1963, p<0.001; Figure 3.d). Furthermore, when analysed separately by the gender, this age-related decline in HERs amplitude remained significant for both male (r = -0.2161; p<0.001) and female (r = -0.1735; p< 0.01). Refer to supplementary tables S3 for specifications of the regression model.

### Relationship of cardiac rhythms and ongoing neural oscillations

To substantiate the hypothesis that cardiac rhythms induce patterned changes in neural oscillations via phase resetting rather than a mere additive enhancement in spectral power (Park et al., 2018), the measure of inter-trial phase coherence (ITPC, see methods section for details) was employed, which increases if phase resetting occurs. Our findings indicate a significant increase in ITPC during the time interval of 180-320ms following the R peak across all age groups (Figure 4.a-c: critical value of ITPC computed at significance level of p= 0.05). However, no time-locked activity was observed ITPC was examined relative to the shuffled ECG time series (preserving the physiological IBI) (Figure 4.d-f). Furthermore, predominant resetting of phase was observed within theta frequency band among young and middle age-group participants.

**Figure 4:**
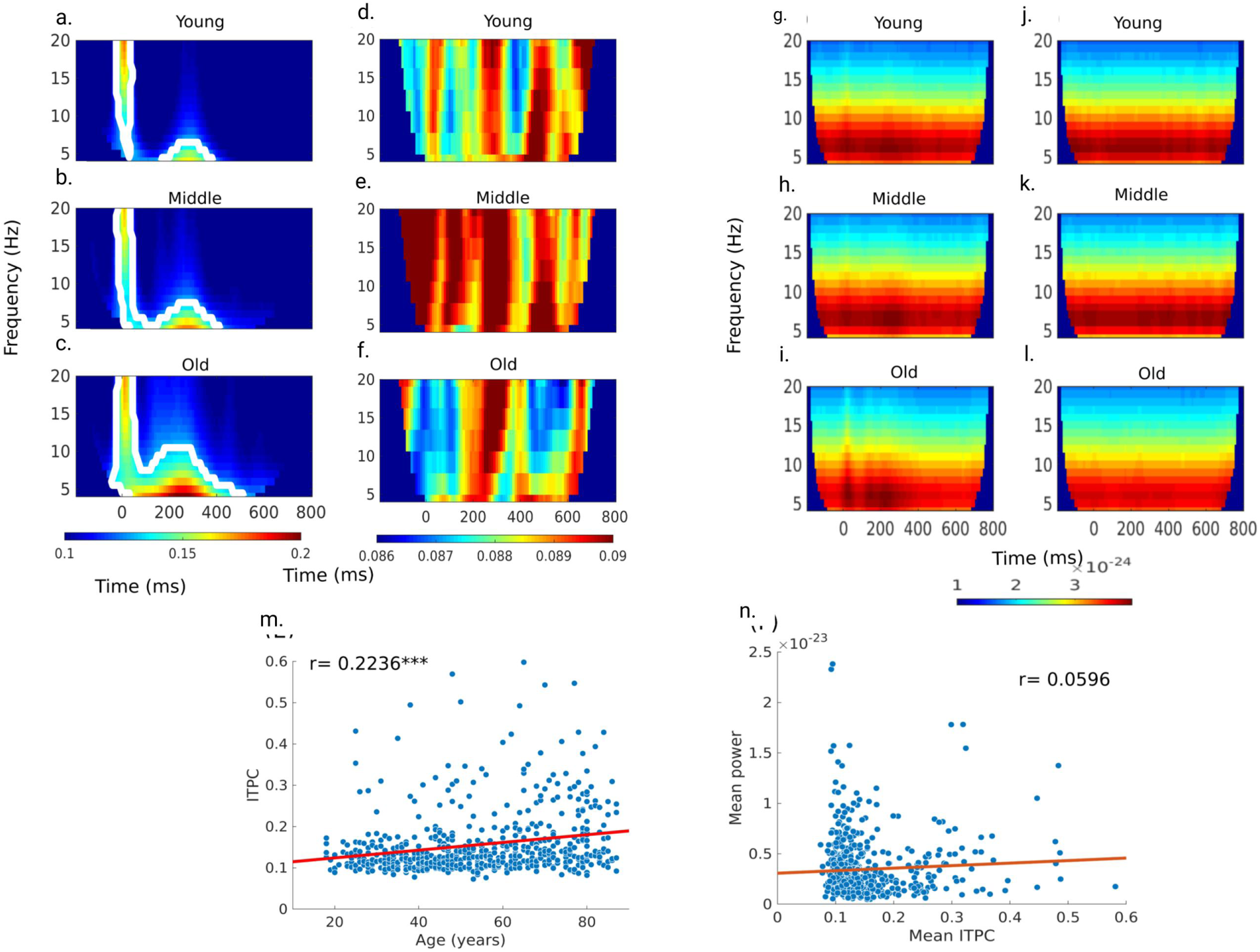
Mechanism of Heartbeat Evoked Responses. Inter trial phase coherence (ITPC) was computed on evoked data time locked to a-c. R peak d-f. shuffled R peak of ECG waveform calculated across all the age group. ITPC values within the white boundary are considered to be statistically significant, using critical value at p = 0.05. Time frequency spectrograms calculated on evoked data time locked to g-i. R peak j-l. shuffled R peak of ECG waveform (found no significant difference between the age groups; cluster permutation test at p =0.05) m. ITPC value (extracted from each subject) in theta band significantly increases across age (r= 0.2236; p<0.001) n. No correlation was observed between mean ITPC values and mean power extracted at around 180-320ms post R peak of ECG waveform for each participant (r=0.0596; p=0.1382).

However, for the older age group, notable activity was observed in the alpha frequency band in addition to the activity in the theta band. The power analysis in the time-frequency domain showed no significant enhanced synchronization time locked to the R peaks (Figure 4.g-i) either for shuffled R peaks (Figure 4.j-l) of the ECG waveform (two-tailed cluster permutation test at p = 0.05). Interestingly, a significant increase in the ITPC values was observed across age in the theta frequency band (r= 0.2236; p<0.001) (Figure 4.m). No significant correlation was observed between mean ITPC values and mean power extracted at around 180-320ms post R peak of the ECG waveform (supplementary table S5; r = 0.0596; p = 0.1382) from each participant (Figure 4.n). No significant ITPC was observed in beta and gamma band in all age groups.

### Underlying cortical sources of HERs

Exact low-resolution electromagnetic tomography (eLORETA) (see Methods for details of implementation) was employed to estimate the underlying sources contributing to HERs observed at the sensor level within the 180-320ms window post R peak of the ECG time series. After the source power was computed for each participant, a grand averaging of the source power was calculated across in participants within each age group (young, middle-aged and elderly) as a group level analysis. All the brain sources were located in the fronto-temporal network (Table 2). The cohort of 620 participants were further divided into the bins of 5 years (18-23, 24-28,29-33,34-38, 39-43, 44-48, 49-53, 54-58, 59-63, 64-68, 69-73, 74-78, 79-83, 84-88 years) for the continuous analysis (Supplementary Table S6). These sources obtained from the continuous analysis were further used to conduct the directed connectivity. Source power surpassing the 95th percentile threshold was considered as significant sources of the HERs in the resting state condition for each age group. The sources obtained by both categorical and continuous analysis was plotted on the glass brain using MNE-Python ^1^. Additionally, all significant sources identified through the continuous analysis were aggregated to compute Granger causality.

**Table 1:** Anatomical labels (according to AAL parcellation) of cortical sources underlying HERs across lifespan.

| Young<br>(N=177) | Middle<br>(N=198) | Old<br>(N=298) |
| --- | --- | --- |
| Right Superior Orbitofrontal Gyrus | Right Superior Orbitofrontal Gyrus | Left Superior Temporal Gyrus |
| Right Middle Orbitofrontal Gyrus | Right Middle Orbitofrontal Gyrus | Right Superior Temporal Gyrus |
| Left Inferior Frontal Gyrus | Right Middle Temporal Gyrus | Left Middle Temporal Pole |
| Right Middle Orbitofrontal Gyrus | Right Middle Orbitofrontal Gyrus | Right Middle Temporal Pole |

**Table 2a:**
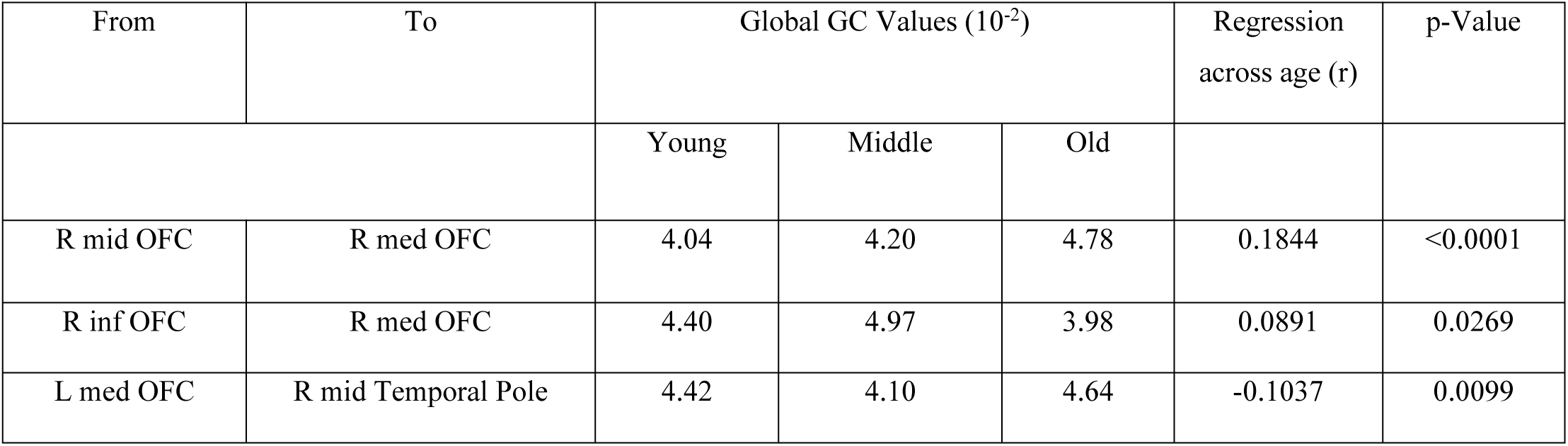
Pairwise list of causally interacting cortical source pairs, showing age-related significant variations in causal strengths (GC values) in different age groups (categorical analysis) and regression coefficients (continuous analysis)

**Table 2b:** Pairwise list of causally interacting cortico-cardiac pairs, showing age-related significant variations in causal strengths (GC values) in different age groups (categorical analysis) and regression coefficients (continuous analysis)

| From | To | Global GC Values ($10^{-2}$) | | | Regression across age (r) | p-value |
| --- | --- | --- | --- | --- | --- | --- |
|  |  | Young | Middle | Old |  |  |
| Cardiac | R superior OFC | 6.26 | 5.51 | 7.2 | 0.1181 | 0.0033 |
|  | L inferior Frontal Gyrus | 6.41 | 5.25 | 5.54 | -0.1099 | 0.0063 |
|  | R middle OFC | 5.95 | 5.87 | 7.51 | 0.1637 | <0.001 |
|  | R superior Frontal Gyrus | 5.39 | 5.59 | 6.54 | 0.1377 | 0.0006 |
|  | R Anterior Cingulate Gyrus | 5.70 | 5.75 | 7.34 | 0.1701 | <0.001 |
|  | R superior OFC | 6.13 | 5.50 | 6.70 | 0.0978 | 0.0151 |
| R superior Temporal Pole | Cardiac | 4.94 | 4.70 | 4.06 | -0.1212 | 0.0026 |
| R superior OFC |  | 4.73 | 5.25 | 4.04 | -0.0915 | 0.0230 |

### Directed functional connectivity between cortico-cortical and cortico-cardiac nodes

Conditional Multivariate Granger causality was computed across all possible HER sources and heart to assess source-level directed connectivity, utilizing the analysis pipeline using Figure 6.a. This analysis provided insights into age-related changes in network dynamics. Heatmaps in Figure 6.c. show GC values between all possible source pairs which are highlighted as nodes in figure 6.b. Figure 6.d. shows the connectivity between these nodes, depicting the strength of GC across different network connections. Notably, several nodes exhibited changes in GC patterns across age, as shown in figure 6.e. Additionally, figure 6.f. demonstrates the connectivity between the heart and brain. Significant causal flow was summarized in Table 3a and Table 3b with the corresponding GC values provided for categorical analysis and regression coefficient for variations with age as a continuous variable. There was a clear increase in strength of causal information flow from heart to brain for all HER sources except left IFG. Interestingly, brain to heart interactions decrease over age. For further details on the casual flow transfer, refer to supplementary table S7.

**Figure 5:**
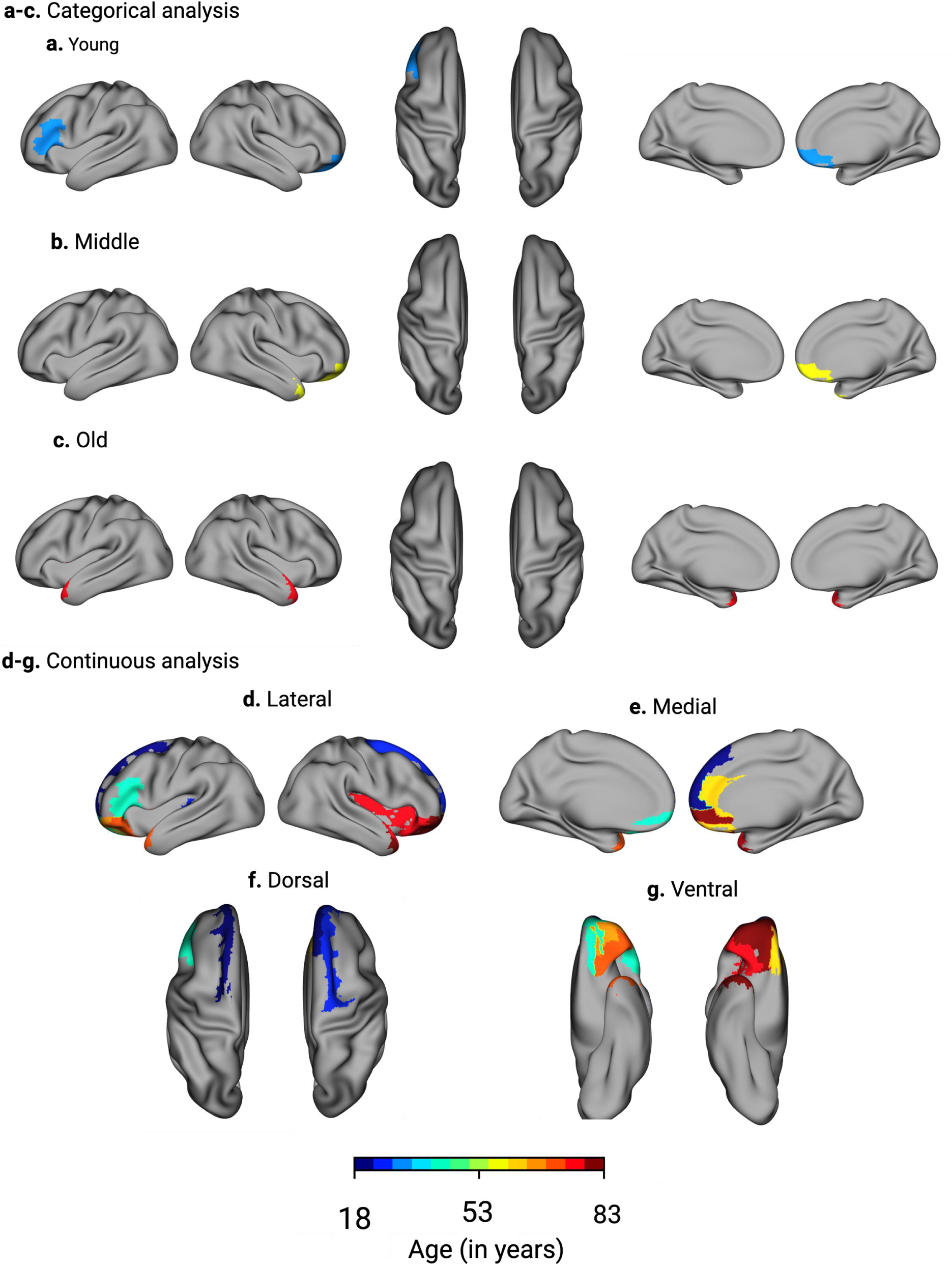
Cortical sources of Heartbeat evoked responses (HERs) across age: Continuous and categorical analysis. The sources represent the cortical areas involved in the processing of heartbeat which includes frontal-temporal regions across age. Panel a-c. represent sources derived from categorical analysis, while panels d-g. show results from continuous analysis. The statistical threshold was set at 95^th^ percentile and the source powers of grid points to surpassing this threshold were considered as significant sources of activation. All regions were approximated to the nearest Broadmann areas of the human brain.

**Figure 6:**
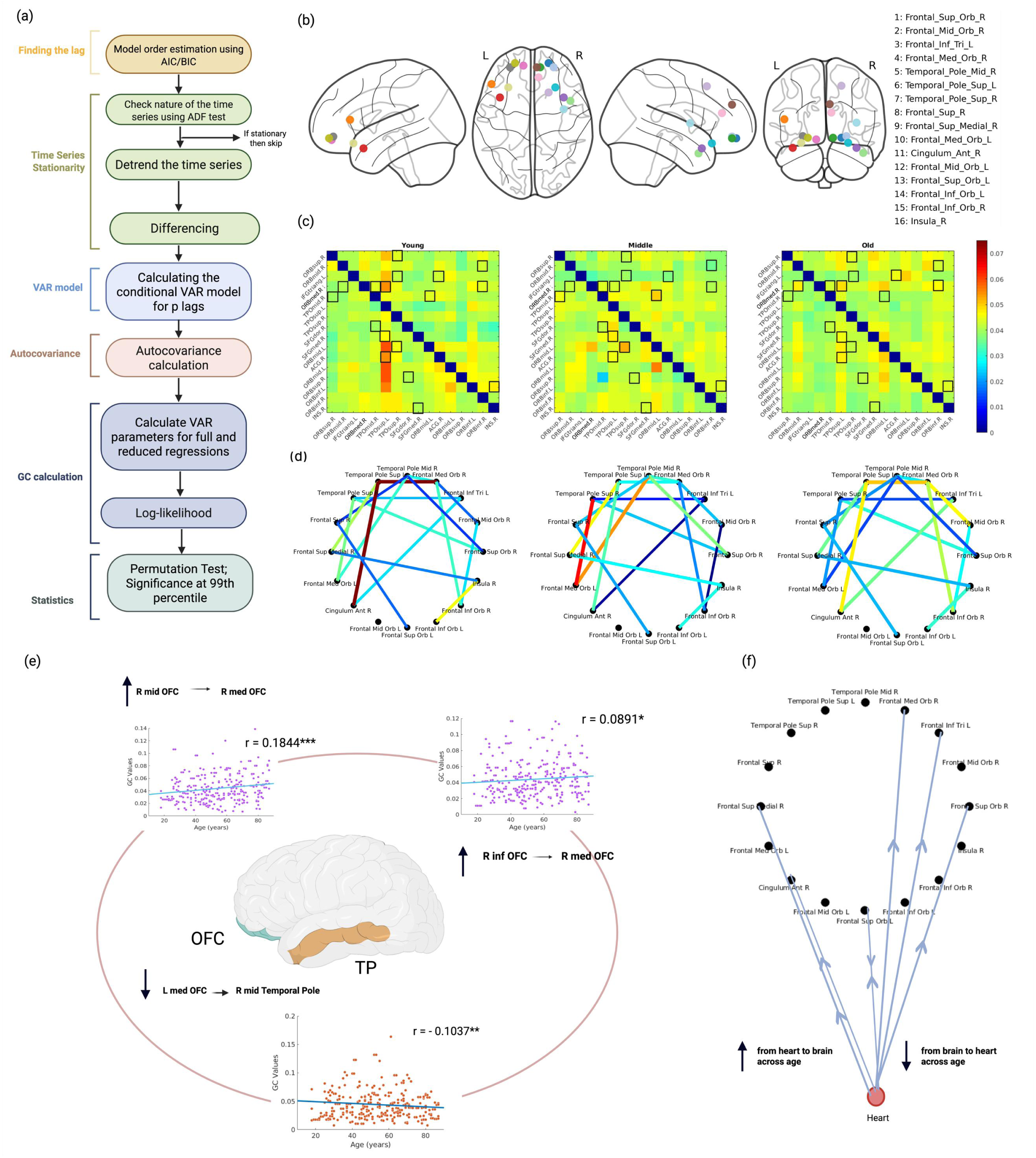
Cortico-cortical and cortico-cardiac information flow across lifespan. a. Workflow used for the calculation of Granger Causality (GC) to evaluate information flow from brain-brain and brain-body interactions b. Visualisation of brain nodes on a glass brain template, used for subsequent cortico-cortical connectivity calcualtion c. GC heatmaps showing connectivity between brain nodes; red indicated high GC value while blue indicates lower GC values. Black boxes highlight statistically significant values after permutation testing followed by Bonferroni correction. d. Connectivity maps displaying significant connection between brain nodes (red line: strong connectivity; blue line: weak connectivity) e. Age related trends in connectivity between and within OFC and TP; connectivity from R mid OFC (r = 0.1846; p<0.0001) and R inf OFC (r = 0.0891; p =0.0269) increases to R med OFC across age; the connectivity from L mid OFC (r = -0.1037;p= 0.0099) to R mid TP decreases across age f. Cortico-cardiac connectivity: connection from the heart to brain increases across lifespan mostly, while connectivity from brain to heart decreases across age.

**Table 3:** Demographic information of Cam-CAN Neuroimaging cohort.

| Measure | Young Adults | Middle Adults | Older Adults |
| --- | --- | --- | --- |
| N | 177 | 198 | 245 |
| Age range (years) | 18-40 | 41-60 | 61-87 |
| Mean age ± standard deviation (years) | 31.32 ± 6.04 | 50.28 ± 5.64 | 72.88 ± 7.27 |
| Female % | 48.02 | 51.01 | 48.39 |

## Discussion

The present study attempted to characterize the age associated changes in the cortical representation of the cardiac signals studied using Heartbeat evoked responses (HERs), neural mechanisms underlying the generation of HERs and corresponding neural networks using a large cross-sectional resting state dataset (N= 620). Four key empirical observations emerged from the subsequent investigations. *First,* time-locked HERs over a temporal window of 180-320 ms was observed post R peak of the ECG waveform (Figure 3) which was consistent for all the participants. HER amplitude in these temporal boundaries decreased across age gradually, establishing the normative patterns of age-related alteration in the processing of cardiac rhythm. *Second*, the mechanism leading to the generation of HER was identified, essentially a phase reset of the spontaneous brain oscillations triggered by the ECG waveform rather than additive spectral power enhancement in theta band (4-8 Hz) frequency band, another alternative that could have been a potential underlying mechanism (Figure 4). Further, a linear increasing trend of the ITPC values across age was observed in the theta (4-8 Hz) frequency band, suggesting that aging is associated with more consistent phase alignment of neural responses to cardiac events and possibly the key reason why perception of time speeds up with age. *Third*, the neural sources of HERs that were identified during the resting state are primarily located in fronto-temporal areas: right superior orbitofrontal, right middle frontal gyrus, left inferior frontal gyrus, right medial orbitofrontal gyrus, right middle temporal pole, left superior temporal pole, right superior temporal pole and left middle temporal pole. Furthermore, a clear pattern of sources shifting from frontal to temporal regions as a function of the age was also identified (Figure 5). *Fourth*, the bidirectional information communication (flow) between the heart and the brain were characterized by advanced time-series analysis to be reorganizing with age; the information flow from the heart to brain majorly increases across age whereas information flow from the brain to heart decreases across age. Along with it, the intraregional connectivity in the orbitofrontal cortex (OFC) increased as a function of age, which might be one of the compensatory mechanisms to account for age related structural decline in large-scale white matter that is observed even during healthy aging (Figure 6). Hence, the present findings firmly quantify the age associated alterations in the cortical representation of cardiac rhythms, from age-related perspective as well as teased out the underlying signal processing mechanism by which such patterns are orchestrated by brain network dynamics which, to be the best of our knowledge, have not been previously reported. In the subsections, we delve into the implications of salient findings from our study.

### Interoceptive abilities decreases as a function of age

What role do HERs play during resting state, why do they undergo age related changes and what implications might these changes have on cognition and emotion? Interoceptive signals, such as heartbeat, act as a metronome for the brain and can be studied through heartbeat-evoked responses (HERs), which reflect cortical responses to cardiac events ^12,15,30^. Here, a time locked neural activity (HER) around 180-320ms post R peak of the ECG waveform and a notable decline in the HER amplitude across age was observed consistently from a categorical and continuous assessment of the data. Prior literature suggests that HER amplitude serves as an index of interoceptive awareness ^31^, which is known to decline across the lifespan ^32^. While there are few speculations for this age-related reduction in interoceptive awareness might be brain undergoing structural changes, such as atrophy in white matter ^33^, degradation of prefrontal cortex ^34,35^, vascular stiffening ^36^ and many more. Bodily signals especially heartbeat, serve not only as the foundation for maintaining homeostasis but also a key role in forming perception of time ^5,9,37^. Enhanced interoceptive accuracy i.e., the ability to perceive one’s heartbeat has been linked to greater temporal precision in sub-second range ^38^. With age, this mechanism may weaken, potentially explaining why the subjective perception of time often appears to accelerate ^39^. Such changes in the temporal processing are likely to impact the broader cognitive, emotional process^39,40^ and disrupted in various neurological disorders such as Alzheimer’s and Parkinson ^2,3^.

### Cardiac signal induces phase resetting on the spontaneous brain activity rather than the amplitude

How does heartbeat influence the spontaneous brain activity? Prior work ^29^ suggests possible basis for generating HERs, the heartbeat acts like an “internal trigger” to the neural oscillations and works by resetting the phase of the spontaneous brain rhythms, rather than by additive evoked responses furthermore connecting it to bodily self-consciousness. In the present study, a comparable phenomenon was observed: ITPC around 180 ms post R peak of the ECG waveform with no change in the spectral power. On shuffling the R peaks of ECG waveform, no time locked activity was observed. Interestingly, oscillation in theta frequency band for the young and the middle age group where strongly phase reset whereas in older participants, in addition to the theta, alpha band was also affected. Moreover, upon extracting ITPC values participant-wise within the theta band, an increasing trend of ITPC across age was observed. Such consolidation of phase resetting indicates an underlying mechanism of a temporal scaffold for neural processing, supporting a consistent internal reference for timekeeping. In younger adults, weaker coupling in the theta phase may allow for greater variability and adaptive integration of interoceptive signal, enabling more nuanced adjustments of subjective time. In contrast, the stronger phase-locking in the theta phase observed with aging may reduce the dynamic range of temporal representations, biasing perception toward a compressed sense of duration and a subjective sense that time passes more quickly with age. These results lay a groundwork for understanding how age-related changes in brain-heart coupling shape the temporal structure of subjective experience along with cognition as well ^4,41^.

### Fronto-temporal network reconfiguration in the representation of cardiac signals

The current findings further suggest that aging is not only associated with change in flexibility in the temporal alignment of interoceptive signals, but also with a broader reorganisation of the networks supporting their cortical representation. In particular, the continuous analysis revealed the gradual shift in the representation of HERs from frontal to temporal regions. With an advancing age, the prefrontal and orbitofrontal cortices may lose their capacity to flexibly integrate interoceptive information, likely reflecting underlying structural changes in these areas ^33,35^. At the same time, the observed shift away from the orbitofrontal cortex which is increasingly recognized as a critical interoceptive hub alongside the insula ^42^, may not reflect a complete loss rather a reweighing of how cardiac signals are represented across lifespan ^43^. Such configuration could indicate that in older individuals, interoceptive signals become less embedded in executive processes and more grounded in affective and memory-related representations, consistent with a lifetime of accumulated experiences and reduced exposure to novelty ^63^. Taken together, these findings might provide a mechanistic account on how age-related changes in interoceptive network organisation, alongside broader network level shifts, may underlie alterations in the subjective experience of time.

### Age related dynamics of cortico-cortical and cortico-cardiac connectivity

With aging, cortical networks reorganize, and their interaction with cardiac signals is altered as well. Cardiac signals relayed to the central nervous system convey information about bodily well-being and also influence cognition ^45^. In turn, the brain sends regulatory feedback to the heart to maintain homeostasis and adapt to environmental demands. Results from this study demonstrate that this bidirectional communication shifts with age: information transfer from the heart to the brain increases, whereas the reverse flow from the brain to the heart progressively declines. The enhanced bottom up influence may reflect a compensatory mechanism, potentially linked to vascular stiffening, while the reduced top-down influence could result from age-related decline in frontal lobe function which regulates the cardiac dynamics. Consistent with this age-related effects on cardiac parameters such as heart rate variability (HRV) and heart rate (HR) was observed. Additionally, strengthening of the connectivity within orbitofrontal cortex (OFC) was observed, whether this represents a true compensation or merely an adaptive change without functional benefit remains an open question.

In summary, the present study demonstrates that aging alters both the cortical representation of cardiac signals and the dynamics of heart-brain communication. HER amplitude decreases while the phase locking increases, reflecting reduced interoceptive awareness and a more rigid temporal scaffolding. Source shifts from frontal to temporal regions and enhanced orbitofrontal connectivity suggest network reorganisation. These alterations are likely to influence time perception, potentially contributing to the well-document sense of accelerated time in older adults. To the best of our knowledge, this is the first study to provide such insights into how cardiac signals shape neural dynamics and providing plausible explanation on subjective time across the lifespan. A key limitation, however, is the absence of the direct interoceptive awareness scores, which could have strengthened the behavioral interpretation of the present findings.

## Methods

### Dataset and Participants

The Cambridge Ageing Neuroscience cohort (Cam-CAN) is a large-scale, multimodal, cross-sectional, population based lifespan (18-88 years) study ^28,46^. The Cam-CAN project involved two stages. During the first stage, cognitive assessments and tests for hearing, vision, balance, and speeded response, were conducted on 2681 participants at their homes. The data collected during stage 1 was also used to screen participants for stage 2. Those with poor hearing, poor vision, neurologic diseases (such as stroke or epilepsy), or a score of less than 25 in the MMSE cognitive assessment examination were excluded from further participation. This study was conducted in accordance with the Helsinki Declaration, and received approval from local ethics committee, Cambridgeshire 2 Research Ethics Committee (reference: 10/H0308/50).

In the second stage of the project, 700 participants (50 men and 50 women from each age band) were screened and recruited to undergo testing at the Medical Research Council (United Kingdom) Cognition and Brain Sciences Unit (MRC-CBSU) in Cambridge. Participants underwent MRI scans, MEG recordings, and completed various psychological tests and neuroimaging assessments during this stage. In the current investigation, only resting-state MEG data was included for analysis, which was acquired concurrently with ECG recordings. Moreover, given that the primary objective of the present study is to examine the cortical representation of the heartbeat across different age groups, the participants were chosen thorough a screening process for cardiovascular disorders and associates disorders. Those who had history of the uncontrolled high blood pressure, stroke or undergone any heart operation or used any blood vessel device/pacemaker were subsequently excluded from the study cohort.

In the present investigation, continuous and categorical approaches were employed to comprehensively capture age-related functional differences. For the continuous analysis, each participant’s age was treated as a single data point. Conversely, for the categorical analysis, the participants were segregated into three distinct groups based on their age: Young adults (18-40 years), Middle adults (41-60 years) and Older adults (61-87 years) as outlined in Table 3.

### Data Acquisition

#### MEG Resting State

The MEG data utilized in this study were obtained from the publicly available Cam-CAN repository, accessible at http://www.mrc-cbu.cam.ac.uk/datasets/camcan/. For all the participants, MEG data were acquired by Elekta Neuromag, Helsinki, using a total of 306 channels, including 102 magnetometers and 204 orthogonal planar gradiometers. The data was collected in a magnetically shielded room (MSR) with low lighting. Data sampling was performed at 1000 Hz with a high-pass filter of 0.03 Hz. Head-position indicator (HPI) coils were used to continuously monitor the position of the participant’s head within the MEG helmet. Additional electrodes were used to record horizontal and vertical electrooculogram (EOG) signals to monitor eye blinks and movements. The acquisition of eyes-closed resting-state MEG data was carried out for at least 8 minutes and 40 seconds.

#### ECG Resting State

ECG data were recorded concurrently with MEG acquisition using a pair of bipolar electrodes with a sampling rate of 1000 Hz. The acquisition lasted at least 8 minutes and 40 seconds.

### Data Preprocessing

The Cam-CAN provided the MEG processed data, which underwent a pre-processing pipeline that involved temporal signal space separation to eliminate noise from HPI coils, environmental sources and continuous head motion correction. A max filter was used to remove the main frequency noise (50-Hz notch filter) and reconstruct any noisy channel. Further information on data acquisition and preprocessing can be found at (Cam-CAN et al., 2014; Taylor et al., 2017).

The MEG time series was detrended, and a bandpass filter was applied with a 1-45 Hz range. Further, ICA (Independent Component Analysis) was employed in MNE-Python to eliminate artifacts such as eye blink and cardiac field.

The ECG time series was detrended before passing it through the Pan-Tompkins algorithm (described in 2.3.1 section) to locate R-peaks in the ECG waveform. Participants were further screened on the basis of their quality of ECG signal (Supplementary Table S8). ECG signals were used to calculate parameters like variability of R-R interval, relative power of the QRS complex, skewness and kurtosis (Supplementary Figure S1). Following the screening, the analysis pipeline on could be performed on the 620 participants (Supplementary Table S1).

### Data Analysis

All the data analysis was conducted in MATLAB using custom-made scripts. First, the fieldtrip toolbox ( https://www.fieldtriptoolbox.org/ ) was used to load the data in the ‘.fif’ format (Oostenveld et al., 2010). The time series corresponds to the 309 x T where 309 are the total number of channels consisting of 102 magnetometers, 204 gradiometers, EOG, ECG and other auxiliary electrodes. For the present study, 102 magnetometers were utilized from the MEG setup and a bipolar ECG electrode. The analysis pipeline is provided in Figure 1.

### Pan-Tompkins Algorithm

The cardiac cycle consists of distinct peaks of P wave, QRS complex and T wave representing mechanical events in the cardiac muscles such as atrial contraction, ventricular contraction and ventricular relaxation respectively. Pan-Tompkins algorithm was employed to locate the QRS complex, the most prominent peak in the ECG waveform (Sedghamiz, 2014). The vector representing the time points of the ECG waveform was passed into the algorithm which involves several steps including bandpass filtering the ECG time series generally in the range of 5-15 Hz, squaring the time series, applying a derivative filter and finally identifying the location of QRS complex using the moving window approach. The algorithm can be found at (https://in.mathworks.com/matlabcentral/fileexchange/45840-complete-pan-tompkins-implementation-ecg-qrs-detector).

### Cardiovascular Scores

The inter-beat interval (IBI) and heart rate variability (HRV) were determined by analyzing ECG signals. The R peaks, identified using the Pan-Tompkins algorithm (refer to section 2.5), were used to extract the IBI, representing the time difference between consecutive R peaks. The average IBI obtained was further used to calculate the heart rate (HR) - the number of heartbeats per unit time (beats/min). HRV was computed by calculating the standard deviation of the IBI, which represents the fluctuations in the time interval between consecutive heartbeats. IBI, HRV and HR were computed for each participant for an initial 60-seconds ECG recording (a short sampling window approach) (Lehavi et al., 2019). For each participant, cam-CAN recorded average systolic and diastolic pressure (recorded three times). To investigate the relationship between these cardiovascular scores and age, linear regression analysis was performed.

### Heartbeat Evoked Responses (HERs)

To map the brain response time-locked to QRS peaks of ECG, the QRS peaks obtained from the Pan-Tompkins algorithm, were used as a trigger points to epoch the MEG time series. This allowed to test the key hypothesis of whether interoceptive signals acts as the stimulant to the ongoing brain dynamics. Based on visual inspection, a pre-stimulus period of 200ms (prior to QRS peak) and a post-stimulus of 800ms (post to QRS peak) was identified that could capture uniquely one complete cardiac cycle. HERs were computed using averaging across trials and channels.

For continuous analysis, the aforementioned HERs was averaged across the number of channels and epochs resulting in a 1 x 1000 dimensional vector where 1000 represents the time points. The amplitude values between 180-320ms post R peak were extracted per subject. To normalize the data, peak-to-peak subtraction was performed by subtracting the maximum amplitude value from each amplitude value for each participant. For categorical analysis, the matrix was stacked according to the age group i.e., younger adults, middle adults, and older adults. The global HERs were obtained by averaging the matrix across channels and trials.

A surrogate analysis as performed to validate the presence of HERs obtained from time locking to R peak of the ECG waveform. This involved computing a shuffled HERs subject wise by randomly selecting non-R peaks to time lock essentially, random points from ECG time series excluding the R peaks, while maintaining the physiological IBI.

### Assessment of the mechanism underlying HERs

**a. Inter-trial phase coherence (ITPC)**

ITPC analysis (Oostenveld et al., 2010) was performed on both the HERs time series synchronized with the R peak and the shuffled HERs time series time-locked to random points in ECG. Power and phases were calculated using wavelet transformation in the frequency range of 4 to 20 Hz for all 102 magnetometers. ITPC was implemented using *ft_freqanalysis* in MATLAB (Oostenveld et al., 2010). The ITPC values are computed using:

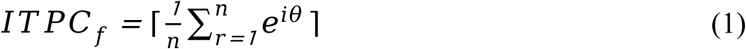

wherein *f* derived is the frequency range at which the ITPC was calculated, 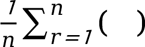 represents an average of complex vector over n trials; *e*^10^ is the Euler’s formula where θ is phase angle derived from complex Fourier coefficients. The value of ITPC ranges between 0 and 1, with higher values of ITPC corresponding to higher phase consistency across trials. As ITPC values exhibit significant variations based on the number of trials or participants, for group analysis consistency, randomly 177 participants were selected (N of the lowly populated group, Table 3) within each group ^47^. To evaluate the significance of phase clustering, the ITPC values obtained were contrasted with a critical value, computed using a significance level of p = 0.05. The critical threshold associated with the p-value was determined as per the formula :

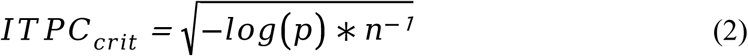

where n represents the total count of trials. ITPC values surpassing this threshold were regarded as statistically significant.

**b. Time frequency estimation**

To delve deeper into understanding the generation of HERs, the power distribution of HERs time series time-locked to R peaks and shuffled HERs time locked to random peaks were examined. The time-frequency analysis was performed by utilizing wavelet transformation with the *ft_freqanalysis* function from fieldtrip toolbox in MATLAB ^48^. Furthermore, cluster permutation test was employed to find the significant clusters from the time frequency analysis, obtained from two conditions i.e., time locked to R peak and activity locked to shuffled R peak using custom built MATLAB code. 1000 iterations of trial randomization were carried out to generate the permutation distribution at every frequency. A two tailed test with a threshold of 0.05 was used to evaluate the time points that exhibit a significant difference in power.

In section 2.8, all the analysis was performed for the 4 to 20 Hz frequency range. This was motivated by earlier findings ^49^ reporting the presence of pulsating artifacts in frequencies below 4 Hz, often attributed to pulsating blood vessels. Our analysis was restricted to the 4-20 Hz frequency range to avoid potential artifacts.

### Source localization of HERs

Source localization was performed to obtain the generators of HERs in the brain averaged across all the participants using a current density technique known as exact low-resolution brain electromagnetic tomography (eLORETA), which was implemented using the MATLAB-based fieldtrip toolbox (Oostenveld et al., 2010). The origin (0,0,0) of the colin27 (Holmes et al., 1998) atlas was set to the anterior commissure and fiducials were set accordingly before generating the head model. Using the single shell method (commonly used for MEG), the brain was segmented into a mesh/grid and employing channel position, the leadfield matrix was computed using *ft_prepare_leadfield.* The regularization parameter (l) was set at 0.1 for the localization of brain sources of HERs. Source power at grid locations (32480 grid points) was calculated using *ft_sourceanalysis* on the HERs time series occurring 180-320 ms post R peak for each age group. The HERs activity was normalized against the baseline activity between interval ([-200 -60]) time locked to R peak of the ECG waveform. Source power was thresholded at 95^th^ percentile, and regions exceeding this threshold were identified as significant sources of activation. These activations were then parcellated using the AAL atlas and visualized with MNE-Python. We have localized the sources for both categorical as well as continuous analysis where we have grouped the participants into three categories (18-40, 41-60, 61-88) and bins of 5 years from 18 to 88 years. For subsequent source time series reconstruction and GC analysis, a total 16 brain nodes were considered, combining sources from the continuous analysis.

### Source time series reconstruction

Time series originating from the significant brain sources were reconstructed, participant-by-participant. This was achieved by multiplying the spatial filter generated from the statistically significant grid points for all the 620 participants to the pre-processed HERs time series (channel x time x trials). A consistent headmodel and electrode placement was used across all the participants. Applying the filter to the HERs epoched time series generated three source dipole time series, oriented along the x, y and z axes. Due to the complexity introduced by three dipole orientations, the time series were projected onto the direction of the strongest dipole. This projection was determined by identifying the dominant eigenvector using singular value decomposition (SVD). Further, to simplify the analysis to estimate GC among few nodes, we applied a k-means clustering approach to classify significant grid points into significant clusters. Sources were grouped into 3 clusters. The centroid of each cluster was identified as a brain area using its proximity to the brain anatomical landmark following the MNI atlas. The time series reconstructed trial-wise for each node was the temporal coefficient projected on the first principal component of the dipolar time series obtained from the cluster. The reconstructed time series from each participant was treated as a trial, making a total of 177 trials for the young, 198 trials for the middle and 245 trials for the older age group.

### Granger Causality

The statistical dependencies among the time series obtained at the source level were investigated by computing the directional interaction among the brain nodes of HERs and cardiac signals to understand the age associated changes associated with it. In this pursuit, conditional multivariate Granger Causality (MVGC) analysis was employed to elucidate the direction and magnitude of the causal influence ^25,27^ among corticocortical and cortico-cardiac interactions. In its most basic form, one can say that any random process, X, Granger causes another random process, Y, when X’s historical occurrences contribute to forecasting Y’s current values to a greater extent than what can be achieved by solely considering Y’s past events while conditioning out additional variables (Z). The GC analysis was conducted using the multivariate granger causality (MVGC) toolbox developed by ^26^. The dataset consisted of the neural signals obtained as source time series from 16 brain nodes and ECG time series (average taken across all the epochs) for each participant. To meet the prerequisites of the GC algorithm, stationarity of these time series was assessed using the Augmented Dickey-Fuller test, employing the *’adftest’* function in MATLAB. Since this revealed non-stationarity in the time series, a difference transformation to induce stationarity was applied, using MATLAB’s *diff* (first order) function. The vector autoregressive (VAR) model was utilized with a lag order of ‘p’, which was estimated by computing Akaike Information Criteria (AIC) using the function ‘*tsdata_to_infocrit’* in the MATLAB based MVGC toolbox (https://in.mathworks.com/matlabcentral/fileexchange/78727-the-multivariate-granger-causality-mvgc-toolbox ). Various values of VAR order, p = 1, 2, ……, 20 was tested. Based on the A/BIC criteria, a VAR order of 6 was chosen for GC estimation. The VAR model is considered robust for the given lag order ’p’ when the Akaike Information Criterion (AIC) and Bayesian Information Criterion (BIC) attain their minimum values. This model can be described as ^26^:

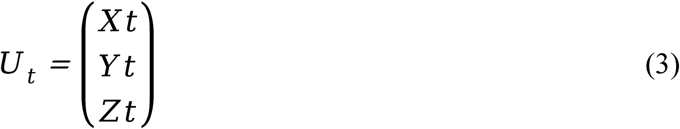

where *U_t_* consists of *X_t_, Y_t_, Z_t_* as the multivariate processes with this idea to eliminate any joint effect of Z on causality from Y to X. The VAR(p) decomposes as into regression coefficient and residuals covariance matrix. From this knowledge, two models i.e., reduced regression and full regression model can be generated and demonstrated as:

Full regression model

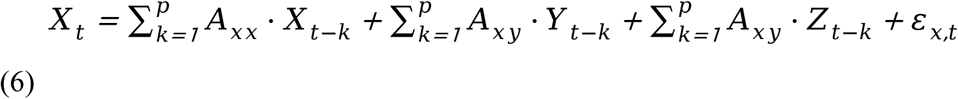

Reduced regression model

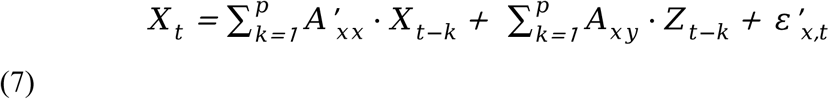

Thus, the Granger causality from Y to X conditioned on Z is calculated by taking the log-likelihood ratio of the covariance residuals obtained from the full and reduced regression model. Mathematically, Granger causality is depicted as:

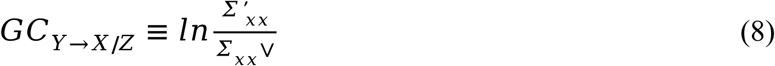

The significance of the GC was calculated against the null distribution generated using the source time series where the phases were staggered while keeping the amplitude conserved. GC peaks in the unshuffled data were considered statistically significant when the observed GC value reached beyond the 99^th^ percentile value of the null distribution. The multiple correction problem was handled by Bonferroni corrections.

### Statistical Analysis

**a. Effect of age and gender on HERs**

To assess the impact of age on HERs, the time series was divided into three segments based on previous literature ^23^ : an early phase (180-320 ms post R peak), a middle phase (460-550 ms post R peak), and a late phase (650-750 ms post R peak). The *anovan* function in MATLAB was used for this analysis. A statistically significant difference between the age groups in the early phase window was observed as compared to the middle and late phase time windows. Furthermore, the impact of gender was examined along with age on HERs. As a result, the early phase HERs window i.e., 180-320 ms post R peak, was selected for further analysis. Refer to Supplementary Table S3 for details.

**b. Eliminating the confound of cardiac field artifact (CFA)**

Given that the selected time window for HERs analysis encompasses the period of 180-320 ms after the R peak, which may be susceptible to contamination by cardiac field artifacts, measures were taken to address this confounding factor. To mitigate the influence of the cardiac field artifact, the ECG amplitude was extracted specifically within the time window of 180-320 ms following the R peak for each of the 620 participants. Subsequently, a linear regression analysis was performed between the mean ECG amplitude obtained from the 180-320 ms post R peak interval and the HERs obtained from the same interval. This regression analysis aimed to account for the ECG activity as a covariate, effectively eliminating its potential confounding effects on the HERs measurements within the specified time window (Supplementary Table S4).

**c. Linear Regression**

Linear regression analysis was performed separately for various parameters (magnitude obtained from peak-to-peak subtraction of HERs in the time interval of 180 to 320ms post R peak, all the cardiovascular measures, estimated ITPC values, GC estimates) as the dependent variable while keeping age as the independent variable.

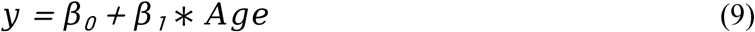

*fitlm.m* function within MATLAB was utilized to perform a linear regression analysis which yielded F-statistics, regression coefficients and goodness of fit *R*^2^ (Supplementary Table S2, S4 ,S6). To assess the strength of the relationship between variables, the correlation coefficient was calculated using the built-in *corrcoef* function available in MATLAB.

## Acknowledgement

The authors acknowledge the generous support of NBRC Core funds and computing facility. This study was supported by NBRC Flagship program BT/MEDIII/NBRC/Flagship/Program/2019: Comparative mapping of common mental disorders (CMD) over lifespan. Data collection and sharing for this project was provided by the Cambridge Centre for Ageing and Neuroscience (Cam-CAN). Cam-CAN funding was provided by the UK Biotechnology and Biological Sciences Research council (grant number BB/H008217/1), together with support from the UK Medical Research Council and University of Cambridge, UK. In accordance with the data usage agreement for Cam-CAN dataset, the article has been submitted as open access.

## Conflict of Interest Statement

All authors declare no conflict of interest.

## Author Credits

**Kirti Saluja:** Conceptualization, Formal analysis, Methods, Investigation, Visualization, Writing-Original draft preparation and editing.

**Dipanjan Roy:** Supervision, Writing-Reviewing and editing.

**Arpan Banerjee:** Conceptualization, Resources, Methods, Investigation, Visualization, Writing-original draft, reviewing and editing, Funding acquisition, Supervision

## Notes

### Competing Interest Statement

The authors have declared no competing interest.

### Summary of Updates

New Analysis on Causal interactions among cortico-cortical and cortico-thalamic sources added A more sharper narrative is used to contextualize the empirical observations

