## Supplemental Material for "When the inner clock fades: Interoceptive decline and consolidation of phase resetting in cortical rhythms by cardiac events underlie healthy lifespan ageing"

The follwing tables support the text and figures presented in the main text.

**Table S1:** Regression table for cardiovascular measures with age. F value, β coefficient, goodness of fit, and significance of the model are reported.

| Independent variable | Dependent variable | F-stats | 1 | $R^2$ | p-value |
|---|---|---|---|---|---|
| Age | Interbeat interval (IBI) | 6.16 | -0.83443 | 0.00987 | 0.0133 |
| | Heartbeat variability (HRV) | 0.00286 | -0.0056268 | 4.63e-06 | 0.957 |
| | Heart Rate (HR) | 7.87 | 0.060119 | 0.0126 | 0.00518 |
| | Systolic Blood Pressure | 125 | 0.40098 | 0.191 | 3.56e-26 |
| | Diastolic Blood Pressure | 2.87 | 0.041845 | 2.87 | 0.091 |

**Table S2**: Effect of age and topography on the Heart evoked responses (HERs) using ANOVA

*3.1. Two Way ANOVA and multiple comparison test applied on 180-320 ms post R peak*

(a)

| Source | SS | df | MS | F | Prob>F |
|---|---|---|---|---|---|
| Columns | 1.44578e-27 | 2 | 7.22888e-28 | 54.87 | 9.30711e-20 |
| Rows | 2.8673e-26 | 101 | 2.83891e-28 | 21.55 | 1.5502e-70 |
| Error | 2.6615e-27 | 202 | 1.31757e-29 | | |
| Total | 3.27803e-26 | 305 | | | |

(b)

| Age group 1 | Age group 2 | Lower Limit | Estimate | Upper Limit | p-value |
|---|---|---|---|---|---|
| Young | Middle | -2.4024e-15 | -1.2111e-15 | -1.985e-17 | 0.045258 |
| Young | Old | -6.2869e-15 | -5.0957e-15 | -3.9044e-15 | 9.5606e-10 |
| Middle | Old | -5.0758e-15 | -3.8846e-15 | -2.6933e-15 | 9.5611e-10 |

*3.2. Two Way ANOVA and multiple comparison test applied on 460-550 ms post R peak*

(a)

| Source | SS | df | MS | F | Prob>F |
|---|---|---|---|---|---|
| Columns | 4.55773e-29 | 2 | 2.27886e-29 | 1.55 | 0.2146 |
| Rows | 6.92583e-27 | 101 | 6.85736e-29 | 4.67 | 0 |
| Error | 2.96852e-27 | 202 | 1.46957e-29 | | |
| Total | 9.94003e-27 | 305 | | | |

(b)

| Age group 1 | Age group 2 | Lower Limit | Estimate | Upper Limit | p-value |
|---|---|---|---|---|---|
| Young | Middle | -1.8179e-15 | -5.5983e-16 | 6.9826-16 | 0.54975 |
| Young | Old | -2.1977e-15 | -9.3961e-16 | 3.1848e-16 | 0.18661 |
| Middle | Old | -1.6379e-15 | -3.7978e-16 | 8.7831e-16 | 0.75906 |

*3.3. Two Way ANOVA and multiple comparison test applied on 650-750 ms post R peak*

(a)

| Source | SS | df | MS | F | Prob>F |
|---|---|---|---|---|---|
| Columns | 5.25924e-30 | 2 | 2.62862e-30 | 1.76 | 0.1748 |
| Rows | 4.13579e-27 | 101 | 4.09484e-29 | 27.39 | 0 |
| Error | 3.0198e-28 | 202 | 1.49495e-30 | | |
| Total | 4.44303e-27 | 305 | | | |

(b)

| Age group 1 | Age group 2 | Lower Limit | Estimate | Upper Limit | p-value |
|---|---|---|---|---|---|
| Young | Middle | -1.2331e-16 | 2.7795e-16 | 6.7922e-16 | 0.23568 |

| | | | | | |
|---|---|---|---|---|---|
| Young | Old | -4.0157e-16 | -3.0375e-19 | 4.0096e-16 | 1 |
| Middle | Old | -6.7952e-16 | -2.7826e-16 | 1.2301e-16 | 0.23494 |

**Table S3**: Regression table for Heart evoked responses with age. F value, β coefficient, goodness of fit, and significance of the model are reported.

| Independent variable | Dependent variable | F-stats | 1 | $R^2$ | p-value |
|---|---|---|---|---|---|
| Age | Heart evoked responses (HERs) amplitude | 24.8 | -1.0356e-16 | 0.0386 | $8.34e^{-07}$ |

**Table S4**: Regression table for ECG amplitude with HERs amplitude extracted from a time window of 180-320 ms post R peak. F value, β coefficient, goodness of fit, and significance of the model are reported.

| Independent variable | Dependent variable | F-stats | 1 | $R^2$ | p-value |
|---|---|---|---|---|---|
| ECG amplitude | Heart evoked responses (HERs) amplitude | 3.19 | -5.1195e-12 | 0.00513 | 0.0746 |

**Table S5:** Regression table for ITPC values and spectral power (conducted for theta frequency band). The ITPC and spectral power values extracted from the significant window (180-400 ms post R peak) and theta frequency range (4-7 Hz). F value, β coefficient, goodness of fit, and significance of the model are reported.

| Independent variable | Dependent variable | F-stats | 1 | $R^2$ | p-value |
|---|---|---|---|---|---|
| ITPC values | Spectral power values | 2.2 | 0.059607 | 0.00355 | 0.138 |

**Table S6:** Cortical sources of Heartbeat evoked responses (HERs) across age : Continuous Analysis

| S.No. | Age group | N | Brain Region |
|---|---|---|---|
| 1 | 18-23 | 20 | Frontal_Sup_R<br>Frontal_Sup_Orb_R<br>Frontal_Sup_Medial_R<br>Frontal_Med_Orb_R<br>Cingulum_Ant_R<br>Thalamus_L |
| 2 | 24-28 | 39 | Frontal_Sup_R<br>Frontal_Med_Orb_R<br>Frontal_Med_Orb_L<br>Rectus_R<br>Cingulum_Ant_R<br>Heschl_L |
| 3 | 29-33 | 41 | Frontal_Sup_Orb_R<br>Frontal_Mid_Orb_L<br>Frontal_Inf_Tri_L<br>Frontal_Med_Orb_L<br>Frontal_Med_Orb_R<br>Cingulum_Ant_R |
| 4 | 34-38 | 55 | Frontal_Sup_Orb_R<br>Frontal_Mid_Orb_R<br>Frontal_Med_Orb_R<br>Rectus_R<br>Amygdala_R<br>Putamen_R |
| 5 | 39-43 | 47 | Frontal_Sup_Orb_L<br>Frontal_Sup_Orb_R<br>Frontal_Mid_Orb_L<br>Frontal_Inf_Tri_L<br>Frontal_Inf_Orb_L<br>Frontal_Med_Orb_L |
| 6 | 44-48 | 62 | Frontal_Sup_Orb_R<br>Frontal_Mid_Orb_R<br>Frontal_Med_Orb_R<br>Rectus_R<br>Cingulum_Ant_R<br>Temporal_Pole_Mid_R |
| 7 | 49-53 | 46 | Hippocampus_R<br>ParaHippocampal_R<br>Amygdala_R<br>Temporal_Pole_Mid_R<br>Cerebullum_3_R<br>Cerebullum_10_R |
| 8 | 54-58 | 53 | Frontal_Mid_Orb_L<br>Frontal_Inf_Orb_L<br>Temporal_Pole_Sup_L<br>Temporal_Pole_Mid_L<br>Temporal_Pole_Mid_R<br>Cerebellum_10_R |
| 9 | 59-63 | 48 | Frontal_Sup_Orb_R<br>Frontal_Mid_Orb_L<br>Frontal_Mid_Orb_R |

| | | | |
|---|---|---|---|
| | | | Frontal_Med_Orb_R<br>Rectus_R<br>Cingulum_Ant_R |
| 10 | 64-68 | 54 | Frontal_Mid_Orb_L<br>Amygdala_L<br>Amygdala_R<br>Putamen_L<br>Putamen_R<br>Temporal_Pole_Mid_R |
| 11 | 69-73 | 47 | Frontal_Inf_Orb_L<br>Olfactory_L<br>Amygdala_L<br>Putamen_L<br>Temporal_Pole_Sup_L<br>Temporal_Pole_Mid_L |
| 12 | 74-78 | 42 | Frontal_Inf_Orb_R<br>Insula_R<br>Putamen_R<br>Pallidum_R<br>Temporal_Pole_Sup_R<br>Temporal_Pole_Mid_R |
| 13 | 79-83 | 54 | Frontal_Sup_Orb_R<br>Frontal_Mid_Orb_R<br>Frontal_Med_Orb_R<br>Rectus_R<br>Amygdala_R<br>Temporal_Pole_Mid_R |
| 14 | 84-88 | 19 | Vermis_3<br>Vermis_4_5<br>Vermis_6<br>Vermis_7<br>Vermis_8<br>Vermis_10 |

**Table S7:** Pairwise list of casually interacting cortico-cortical and cortico-cardiac pairs

*(a) Significant causal interaction between the cortico-cortical nodes*

| From | To | Global GC values ($10^{-2}$) |
|---|---|---|
| Frontal_Sup_Orb_R | Temporal_Pole_Mid_R | 4.16 |
| Frontal_Mid_Orb_R | Frontal_Med_Orb_R | 4.38 |
| Temporal_Pole_Mid_R | Frontal_Med_Orb_R | 4.34 |
| Temporal_Pole_Mid_R | Frontal_Sup_R | 4.10 |
| Temporal_Pole_Sup_L | Frontal_Med_Orb_R | 4.87 |
| Temporal_Pole_Sup_L | Frontal_Sup_Med_R | 4.62 |
| Temporal_Pole_Sup_L | Cingulum_Ant_R | 4.88 |
| Temporal_Pole_Sup_R | Frontal_Sup_Orb_R | 4.33 |
| Temporal_Pole_Sup_R | Frontal_Inf_Tri_L | 4.01 |
| Temporal_Pole_Sup_R | Frontal_Med_Orb_L | 4.60 |
| Frontal_Sup_R | Frontal_Sup_Orb_L | 4.13 |
| Frontal_Sup_Med_R | Insula_R | 4.20 |
| Frontal_Med_Orb_L | Temporal_Pole_Mid_R | 4.42 |
| Cingulum_Ant_R | Frontal_Inf_Tri_L | 4.18 |
| Frontal_Inf_Orb_R | Frontal_Mid_Orb_R | 4.08 |
| Frontal_Inf_Orb_R | Frontal_Med_Orb_R | 4.40 |
| Insula_R | Frontal_Inf_Orb_L | 4.49 |

*(b) Significant causal interaction between the cortico-cardiac nodes*

| From | To | Global GC values ($10^{-2}$) |
|---|---|---|
| Frontal_Inf_Tri_L | Cardiac | 4.79 |
| Temporal_Pole_Mid_R | | 4.75 |
| Temporal_Pole_Sup_L | | 4.74 |
| Temporal_Pole_Sup_R | | 4.51 |
| Frontal_Sup_R | | 4.74 |
| Frontal_Sup_Med_R | | 4.52 |
| Frontal_Med_Orb_L | | 4.83 |

| | | |
|---|---|---|
| Cingulum_Ant_R | | 4.61 |
| Frontal_Mid_Orb_L | | 5.47 |
| Frontal_Inf_Orb_L | | 5.12 |
| Frontal_Inf_Orb_R | | 4.72 |
| Insula_R | | 4.52 |
| Cardiac | Frontal_Sup_Orb_R | 6.39 |
| | Frontal_Mid_Orb_R | 5.87 |
| | Frontal_Inf_Tri_L | 5.69 |
| | Frontal_Med_Orb_R | 6.54 |
| | Temporal_Pole_Mid_R | 5.90 |
| | Temporal_Pole_Sup_R | 6.36 |
| | Temporal_Pole_Sup_L | 6.33 |
| | Frontal_Sup_R | 5.45 |
| | Frontal_Sup_Medial_R | 5.91 |
| | Frontal_Med_Orb_L | 6.27 |
| | Cingulum_Ant_R | 6.36 |
| | Frontal_Mid_Orb_L | 5.85 |
| | Frontal_Sup_Orb_L | 6.15 |
| | Frontal_Inf_Orb_L | 6.10 |
| | Frontal_Inf_Orb_R | 5.92 |
| | Insula_R | 5.76 |

**Table S8:** List of participants excluded from the study along with their exclusion criteria. The exclusion criteria based on ECG signal processing measure such as skewness, high R-R coefficient of variance, relative QRS complex power
Initial number of participants : **650**
Participants after exclusion : **620**

| S.No. | Subject ID | Age | Exclusion Criteria |
|---|---|---|---|
| 1 | CC310160 | 41 | ECG signal not available |
| 2 | CC420729 | 56 | ECG signal not available |
| 3 | CC610050 | 71 | ECG signal not available |
| 4 | CC610061 | 76 | ECG signal not available |
| 5 | CC620044 | 73 | ECG signal not available |
| 6 | CC620499 | 71 | ECG signal not available |
| 7 | CC620557 | 74 | ECG signal not available |
| 8 | CC710088 | 83 | ECG signal not available |
| 9 | CC710154 | 83 | ECG signal not available |
| 10 | CC711035 | 88 | ECG signal not available |
| 11 | CC720071 | 82 | ECG signal not available |
| 12 | CC720103 | 80 | ECG signal not available |
| 13 | CC721519 | 79 | ECG signal not available |
| 14 | CC722077 | 82 | ECG signal not available |
| 15 | CC110045 | 24 | High R-R coefficient of variance |
| 16 | CC121411 | 26 | High R-R coefficient of variance |
| 17 | CC620152 | 73 | High R-R coefficient of variance |
| 18 | CC720685 | 81 | High R-R coefficient of variance |
| 19 | CC410121 | 52 | High skewness and kurtosis value |
| 20 | CC420061 | 57 | High skewness and kurtosis value |
| 21 | CC722542 | 79 | High skewness and kurtosis value |
| 22 | CC410226 | 56 | Relative power of the QRS complex is high due to a split in the R peak |
| 23 | CC420180 | 56 | Relative power of the QRS complex is high due to a split in the R peak |
| 24 | CC610288 | 69 | Relative power of the QRS complex is high due to a split in the R peak |
| 25 | CC610658 | 78 | Relative power of the QRS complex is high dude to a split in the R peak |
| 26 | CC720646 | 81 | Noisy signal |
| 27 | CC410094 | 54 | Noisy trials (MEG) |

| | | | |
|---|---|---|---|
| 28 | CC510480 | 68 | Noisy trials (MEG) |
| 29 | CC620490 | 74 | Noisy trials (MEG) |
| 30 | CC620436 | 78 | Noisy trials (MEG) |
